# Scene-relative object motion biases depth percepts

**DOI:** 10.1101/2022.06.22.497241

**Authors:** Ranran L. French, Gregory C. DeAngelis

## Abstract

An important function of the visual system is to represent 3D scene structure from a sequence of 2D images projected onto the retinae. During observer translation, the relative image motion of stationary objects at different distances (motion parallax) provides potent depth information. However, if an object moves relative to the scene, this complicates the computation of depth from motion parallax since there will be an additional component of image motion related to scene-relative object motion. To correctly compute depth from motion parallax, only the component of image motion caused by self-motion should be used by the brain. Previous experimental and theoretical work on perception of depth from motion parallax has assumed that objects are stationary in the world. Thus, it is unknown whether perceived depth based on motion parallax is biased by object motion relative to the scene.

Naïve human subjects viewed a virtual 3D scene consisting of a ground plane and stationary background objects, while lateral self-motion was simulated by optic flow. A target object could be either stationary or moving laterally at different velocities, and subjects were asked to judge the depth of the object relative to the plane of fixation. Subjects showed a far bias when object and observer moved in the same direction, and a near bias when object and observer moved in opposite directions. This pattern of biases is expected if subjects confound image motion due to self-motion with that due to scene-relative object motion. These biases were large when the object was viewed monocularly, and were greatly reduced, but not eliminated, when binocular disparity cues were provided. Our findings establish that scene-relative object motion can confound perceptual judgements of depth during self-motion.

## Introduction

Many situations, some critical to an animal’s survival such as hunting prey, require the brain to correctly identify moving objects in a three-dimensional (3D) scene during self-motion. To make sense of the 3D structure of the visual scene, the brain must also accurately infer the depths of moving objects. The visual system has evolved multiple mechanisms to judge the depth of objects in a 3D scene ^1-3^. Stereoscopic depth perception relies on slight differences between the retinal images projected onto the two eyes, known as binocular disparity ^4, 5^. Robust monocular depth perception can also be attained from motion parallax, the image motion induced by translational self-motion ^6-8^. Moreover, the brain can estimate depth with greater precision by integrating disparity and motion parallax cues ^9-11^.

Regarding depth perception from motion parallax, stationary objects at varying depths have different retinal image velocities during lateral self-translation. In Figure 1a, when an observer maintains visual fixation on the traffic light while moving to the right, all of the stationary objects in the scene have an image velocity that is determined by their 3D location and the movement of the observer. After taking into account image inversion due to the lens of the eye, stationary objects that are nearer than the point of fixation (traffic light) would have rightward image motion, whereas stationary objects that are farther than the traffic light would have leftward image motion (Figure 1b,d). In this situation, a stationary object’s depth relative to the fixation plane, *d*, is determined by the retinal image velocity, *dθ*/*dt*, eye rotation velocity relative to the scene, *dα*/*dt*, and the fixation distance, *f* (Figure 1b). Mathematically, relative depth is given by the motion-pursuit law ^12^, which can be approximated for small angles as:

**Figure 1:**
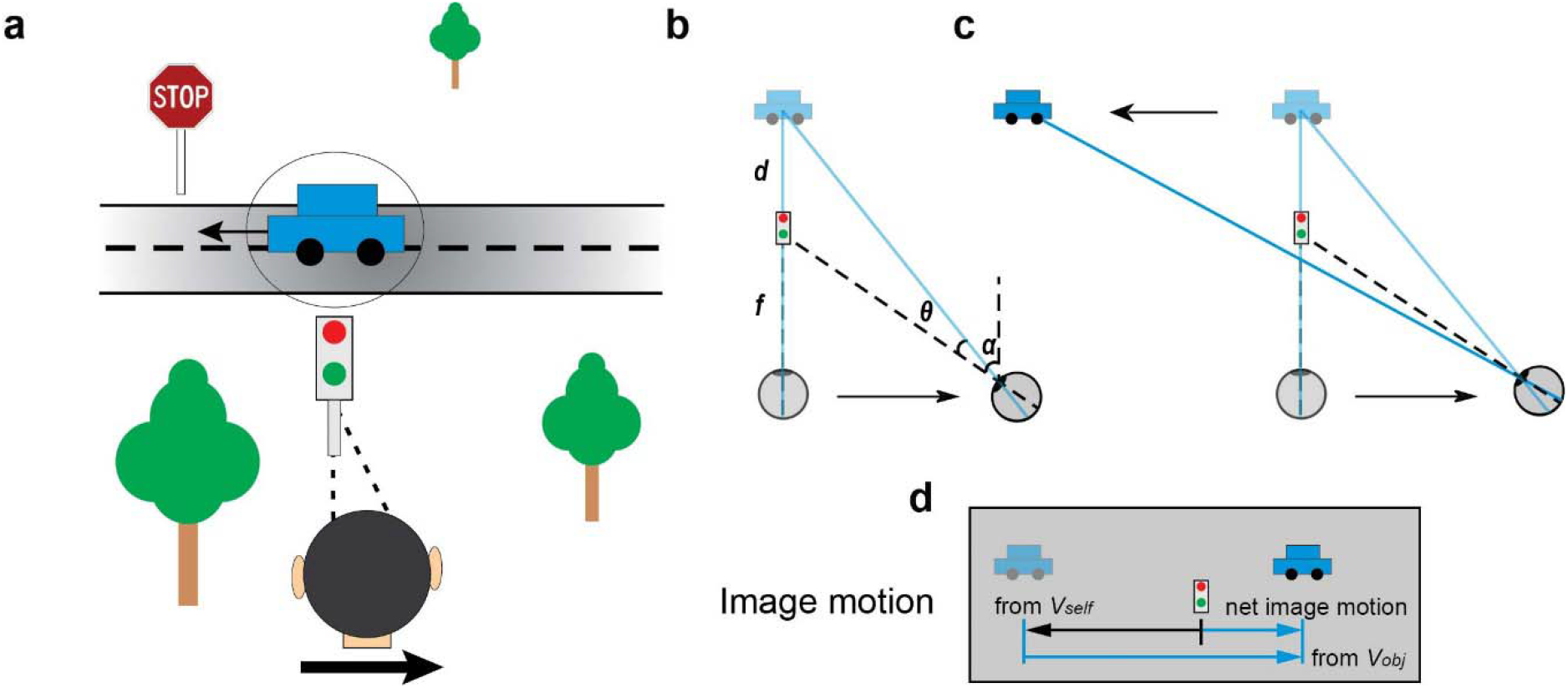
Schematic illustration of motion components that arise from observer translation and scene-relative object motion. a) An observer fixates on the traffic light while translating to the right (adapted from Nadler et al 2013). At the same time, a car moves independently to the left in the scene. b-d) Illustration of the components of image motion of the car related to self-motion and scene-relative object motion. Note that image motion in panel d takes into account image inversion by the lens of the eye. b) If the car is stationary in the world, it has a leftward image motion component due to self-motion, *v*_*self*_ (lighter shading). c) If the car also moves to the left in the world while the observer moves to the right, image motion of the car also has a component resulting from object motion, *v*_*obj*_. d) The net image motion of the car is to the right.

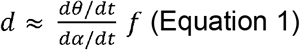

Thus, the relative depth of stationary objects can be computed from their image motion if eye velocity relative to the scene is also known.

Although much has been learned about how humans perceive depth from motion parallax ^7, 13-22^, previous studies have not considered situations in which the object of interest may be moving in the world. Judging depth of a moving object during self-motion is substantially more challenging, as the retinal image velocity, *v*_*ret*_, depends on both scene-relative object motion and self-motion (Figure 1d). For the moving car in Figure 1a, there is a component of image motion related to the car’s motion in the world, *v*_*obj*_, and a component of image motion related to the observer’s translation and the car’s location in depth, *v*_*self*_ (Figure 1d). To correctly compute the depth of the moving car from motion parallax using the motion-pursuit law (Eqn. 1), one would need to use only the component of image motion related to self-motion, *v*_*self*_, while ignoring the component of image motion due to the object’s motion in the world, *v*_*obj*_. In other words, to compute depth accurately via the motion pursuit law, *dθ*/*dt* in Eqn (1) should equal *v*_*self*_. If the observer cannot successfully parse out *v*_*self*_ and incorporates at least some portion of *v*_*obj*_ into *dθ*/*dt*, then the resulting depth estimate would be erroneous.

Figure 2 further illustrates how scene-relative object motion could bias perception of depth based on motion parallax. Figure 2a depicts a physical scenario in which: 1) an object is located nearer than the fixation point, 2) the object moves to the right in the world (magenta circles), and 3) the observer translates to the right (same direction as the object) while counter-rotating their eye to maintain fixation on a world-fixed point (green cross). In this scenario, the net image motion, *v*_*ret*_, (Figure 2g, magenta dots) is also consistent with that produced by a stationary object at a farther depth (Figure 2d, magenta circle). If the subject can correctly infer the true physical scenario (Figure 2a) and can parse out the component of image motion due to self-motion, *v*_*self*_, then depth perception should be unbiased (Figure 2c, magenta curve). However, if the subject incorrectly infers that the object is stationary in the world (Figure 2d) and computes depth based on the net image motion, we expect a substantial far bias (Figure 2f, magenta curve). As typically done in studies of depth from motion parallax ^23^, depth in Figure 2c,f is expressed in units of equivalent binocular disparity. This is done such that near and far objects having the same magnitude of equivalent disparity would produce the same motion parallax resulting from observer translation.

**Figure 2:**
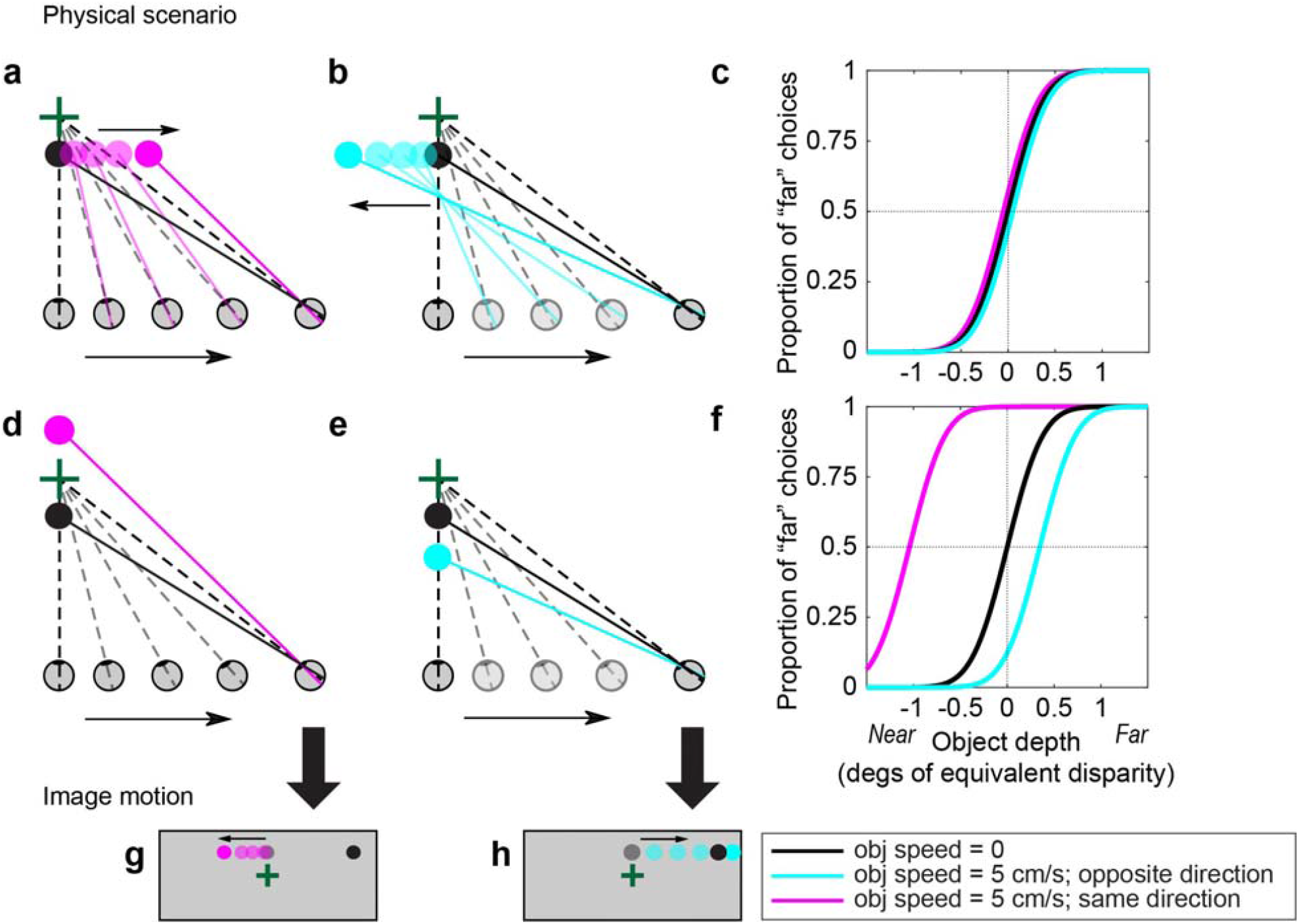
Predicted behavioral results under two hypotheses. a) Example physical scenario in which an object (magenta circles) and the observer’s eye are translating in the same direction (rightward). As the observer translates, they counter-rotate their eye to maintain fixation on a world-fixed point (green cross). The black circle shows the location of the object if it is stationary in the world (obj speed = 0). b) Example scenario in which the object (cyan circles) and self are moving in opposite directions. c) Predicted depth psychometric curves if an observer can accurately parse retinal image motion into components related to object motion and self-motion, and use the latter to compute depth via the motion-pursuit law. Note that depth is expressed in units of equivalent binocular disparity (see text). d) Physical scenario in which a stationary object at a far depth produces the same image motion as in panel a. e) Physical scenario in which a stationary near object creates the same image motion as in panel b. f) Predicted depth psychometric curves if an observe computes depth, via the motion-pursuit law, based on net image motion. Note that the magnitude of the perceptual bias, in degrees of equivalent disparity, is smaller when the object moves in the direction opposite to self-motion (cyan curve). This is due to the nonlinear relationship between depth measured in metric distance versus equivalent disparities. g) Image motion corresponding to panels a and d. h) Image motion corresponding to panels b and e.

Analogously, when an object is moving to the left in the world (Figure 2b, cyan circles), opposite to self-motion, the net image motion could also be consistent with that produced by a stationary object at a nearer depth (Figure 2e, cyan circle). In this case, if the subject attributes the net image motion only to self-motion and thinks that the object is stationary, then we would expect perceived depth to have a near bias (Figure 2f, cyan curve). Thus, by measuring biases in perceived depth due to the addition of scene-relative object motion, we can measure the extent to which human observers are able to discount image motion associated with scene-relative object motion, *v*_*obj*_, and accurately compute depth.

If there were no other cues to depth besides motion parallax, then the prediction of Fig. 2f is expected to hold, as there would be no cues to distinguish *v*_*obj*_ from *v*_*self*_. In our experiments, we include a robust size cue in the monocular viewing condition that could, in principle, allow observers to dissociate *v*_*obj*_ and *v*_*self*_. We also include a binocular condition in which there are powerful binocular disparity cues to depth, and we expect that any biases observed during monocular viewing would be decreased in the presence of binocular disparity cues. In the situations considered here, scene-relative object motion produces a component of image motion that is compatible with (in the same direction as) self-motion (Figures 1, 2). Of course, in the real world, objects may move in ways that makes it much easier to dissociate *v*_*obj*_ from *v*_*self*_ (e.g., a bird flying near vertically in Figure 1a). Thus, our experiment presents a rather stringent test to the visual system.

Our results demonstrate that humans do not completely isolate image motion associated with self-motion and accurately compute depth from motion parallax, at least under conditions in which the image motion associated with scene-relative object motion is also compatible with self-motion. In the monocular condition, for which object size provides the only other depth cue, subjects have very large biases consistent with attributing virtually all image motion to self-motion. In the binocular condition, biases are greatly reduced but not eliminated, indicating that scene-relative object motion can still bias depth perception even in the presence of powerful binocular disparity cues.

## Methods

### Subjects

The experiments were approved by the Institutional Review Board at the University of Rochester (Study ID: 00003094), and all methods were carried out in accordance with institutional guidelines and regulations. A total of 8 healthy adults (17—23 years old) were recruited for this study and none of the subjects was aware of the goal of the study. Two subjects, 227 and 228, were screened out before starting data collection due to low quality data acquired with our eye tracking system, leaving 6 subjects for the main data set. Subjects were informed of the experimental procedures and written informed consent was obtained from each human participant. All subjects had normal or corrected-to-normal vision.

### Setup and visual stimuli

Each subject was seated in front of a 27’’ computer monitor (BENQ, XL2720) which operated at 1920 × 1080 pixel resolution and a 120 Hz refresh rate. Visual stimuli were generated using the OpenGL libraries and rendered using an accelerated graphics card (Nvidia Quadro K4200), with anti-aliasing for smooth motion. The room was darkened for the duration of the task. The viewing distance was 57 cm, such that the display subtended 60 degrees horizontally and 33.6 degrees vertically. The subject’s head was stabilized by a chin and forehead rest. The subject’s eyes were tracked by an Eyelink II (SR-Research) infrared eye tracker. Eye movement signals were calibrated at the beginning of each session and were stored for off-line analyses.

The subject was instructed to fixate their gaze on a fixation point, which was presented at eye height and initially located at the center of the display prior to self-motion. While we monitored eye movements on-line and analyzed them off-line, visual fixation and pursuit tracking were not enforced from trial to trial during task performance. In other words, if a subject broke fixation during a trial, the trial was not aborted and redone. Rather, we accepted behavioral reports for all trials, regardless of eye movements, and we analyzed the relationship between eye movements and task performance post-hoc. The fixation point was rendered stereoscopically to lie at a simulated distance of 3.57 m; that is, 3m farther from the subject than the visual display. The fixation point was a square that subtended 9.4 arcseconds horizontally and vertically. To ensure that subjects’ eyes could diverge properly to fuse the fixation point, we asked them to report the color of the fixation point. Since the stereoscopic scene was viewed through red and green colored filters (Kodak Wratten2 filter nos. 29 and 61) for the left and right eyes, respectively, we considered that the subject had successfully fused the fixation point if they reported seeing it as yellow and did not report seeing separate red and green images. Otherwise, we brought the fixation point to the screen distance, and successively increased the distance of the fixation point in increments of 0.13 degrees of disparity. For each increment, we asked the subject whether the fixation point appeared yellow. If the subject no longer perceived the fixation point as yellow, we would go back one step until their vergence stabilized again.

The visual scene included a ground plane composed of randomly oriented equilateral triangles with each side extending 40 cm in world coordinates. The density of these triangles was 2 triangles/m^2^. The ground plane was visible from 0.05 m to 17.5 m in front of the viewer. The height of the OpenGL camera was 1.25 m above the ground, comparable to most subjects’ eye height when they were seated in front of the monitor. There were also nine scene-fixed (equivalent to world-fixed in our experiment) stationary landmarks whose locations and sizes varied from trial to trial (Figure 3). Each landmark was an ellipsoid composed of randomly-positioned equilateral triangles with density of 0.02 triangles/cm^3^ and each edge of the triangle extending 4 cm in world coordinates. The positions of the nine stationary landmarks were chosen on each trial from within a 3-cell x 3-cell invisible grid, so that the landmarks did not overlap. We define the origin of the world-centered coordinate frame as the location of the cyclopean eye. The invisible grid spanned from -3 m to +3 m along the x-axis and from 1.5 m to 10.5 m along the z-axis. Each cell in the grid was 2 m wide and 3 m deep and each landmark’s position only varied within a cell. The width, height, and depth of the landmarks varied between 50 and 70 cm.

**Figure 3:**
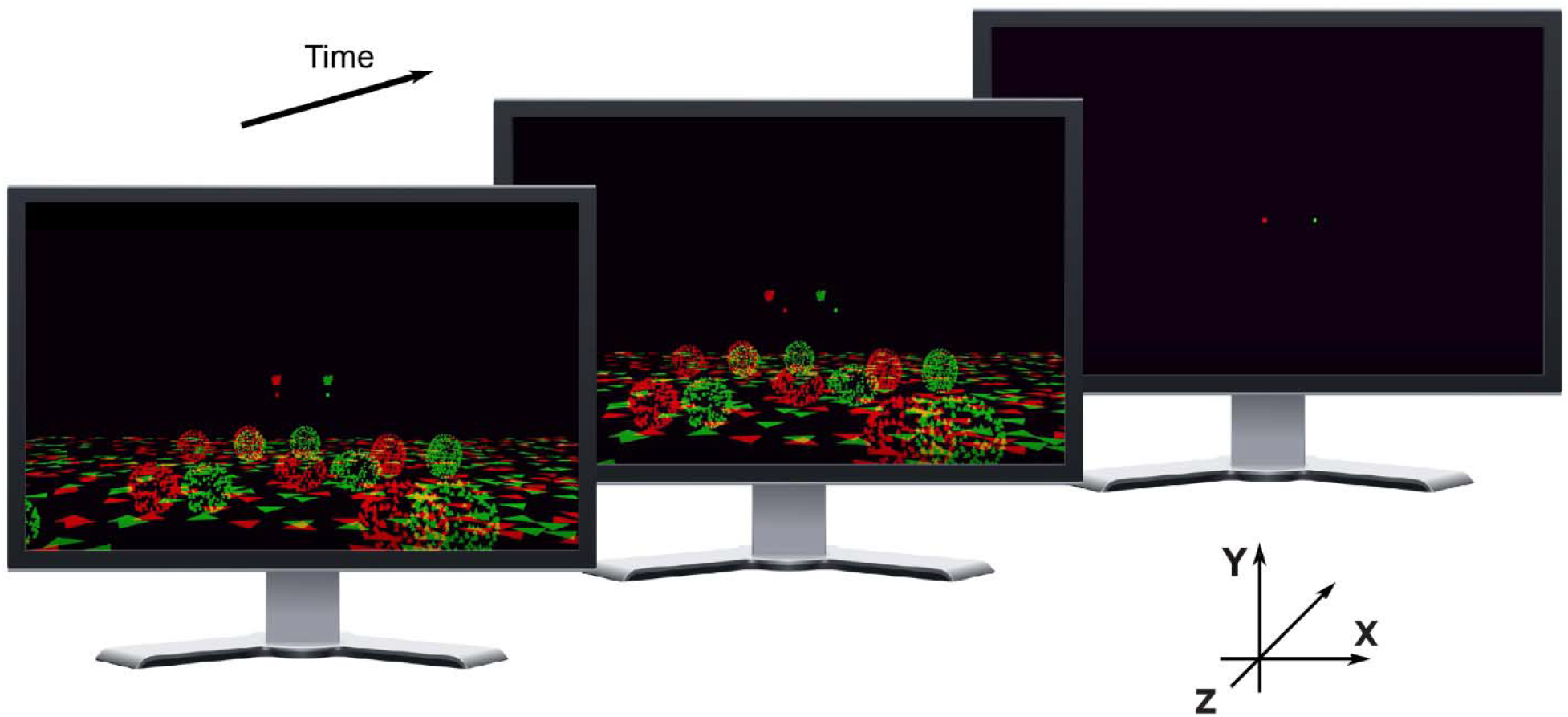
Illustration of visual stimuli used in the psychophysical experiment. The scene consists of a textured ground plane, nine stationary objects of varying size and location that sit on the ground plane, and an object in the upper visual field that may be stationary or moving relative to the scene. Stereo images show a sequence of actual frames of the display for a situation in which the target object is presented binocularly. Red and green elements show images presented to the left and right eyes via red-green anaglyph glasses. The fixation point is always located above the horizon and the object of interest is seen above the fixation point. In the particular stimulus condition illustrated here, the target object lies at approximately the same distance as the fixation point, such that motion of the target relative to the fixation point (middle panel) results predominantly from independent object motion relative to the scene, rather than from self-motion. Illustrations by R.L. French.

Optic flow simulating lateral self-motion was generated by translating the OpenGL cameras (one for each eye) along the x-axis (parallel to the interaural axis). Simulated self-motion had a Gaussian velocity profile with a standard deviation of 0.667 s and a peak velocity at 22 cm/s. The resultant optic flow field contained naturalistic cues that mimicked translation of the observer through the 3D virtual world, including motion parallax, size variations, and binocular disparity.

The object of interest was a small “ball” of dots, which could be viewed either binocularly (through red and green filters) or monocularly (left eye, red filter only). The object was a transparent sphere with a diameter of 8 cm was composed of random dots having a density of 0.05 dots/cm^3^. The size of the dots comprising the object was fixed in pixel units, such that dot size did not change with distance to the object. The object translated only along the horizontal axis (x-axis in Figure 3), and had a fixed vertical location in angular coordinates (1.6 degrees above the fixation point). The vertical location of the object in the world was adjusted as a function of its depth to keep the vertical location fixed in retinal coordinates. The object’s location in depth (z-axis) was varied from trial to trial as described below. The horizontal starting position of the object along the x-axis was randomized following a normal distribution with a mean of 0 and a standard deviation of 1.36 cm, which made the starting and ending positions less correlated with the object’s depth. The object had a fixed physical size such that its image boundary scaled with its location in depth, even though dots within the object maintained a fixed retinal size. Depth values were specified in units of binocular disparity when the stimulus was dichoptic, or in units of equivalent disparity for the monocular condition ^23^. Visual stimuli were presented for 2 s in each trial. See Video 1 for example stimuli.

The subjects were reminded at the beginning of each session to track the fixation point with their eyes, not the target object. We also encouraged subjects to blink between trials as needed.

### Main experiment: design and procedure

Each trial began with presentation of a world-fixed fixation point, and the remainder of the visual stimulus appeared after a delay of 500ms. The stimulus was presented for 2s; however, due to the Gaussian velocity profiles used for both self-motion and object motion, the visual scene was initially stationary on the display and began moving gradually. Subjects were instructed to track the fixation point throughout the duration of the stimulus and to make a judgement about the depth of the object relative to the fixation point. Because the fixation point was fixed in the virtual environment, simulated self-motion caused the fixation point to translate horizontally on the visual display. Thus, to maintain fixation, subjects were required to smoothly rotate their eyes in the direction opposite to simulated self-motion, in order to maintain their gaze on the fixation point. After the visual stimulus disappeared, subjects had up to 1.5 s to report their perceived depth by pressing one of two keys (“near” vs “far”) on a key pad to indicate whether they thought the object was nearer or farther than the fixation point.

All subjects completed a training phase and a testing phase for both binocular and monocular conditions. Each training phase included 70 trials (7 different depths x 2 self-motion velocities x 5 repetitions). During the training phase, the object of interest was always stationary in world coordinates and the subjects were told this. Subjects were also informed that the object of interest maintained a fixed physical size in the virtual environment, such that image size was a cue to depth. In addition, during the training phase, subjects were given auditory feedback in which a high-pitched beep indicated a correct depth judgement and a low-pitched beep indicated an error. Each testing phase included 1050 trials (7 different depths x 5 object velocities x 2 self-motion velocities x 15 repetitions). During the testing phase, the object may or may not have moved relative to the scene, and subjects were told explicitly that the object may or may not move independently in world coordinates. No feedback was provided to subjects during the testing phase. Subjects always underwent binocular training and testing phases first before moving on to the monocular conditions. This sequence helped subjects get used to the task during binocular conditions in which disparity cues greatly aided depth discrimination. Note that, because monocular conditions were run second, the much smaller biases observed in the binocular condition cannot be a result of subjects learning to better parse image motion into its components over time.

The depth ranges were different between binocular and monocular conditions due to the reduced sensitivities of subjects without the stereo cue. The depth values, relative to the fixation point, used in binocular conditions were 0, ±0.02, ±0.04, and ±0.06 degrees; the corresponding depths used in monocular conditions were 0, ±0.25, ±0.5, and ±0.75 degrees. Here, positive depth values correspond to uncrossed disparities associated with far distances, and negative values correspond to crossed disparities associated with near distances. Depths are specified in units of equivalent disparity such that motion parallax resulting from self-motion has the same speed for near and far objects with the same magnitude of equivalent disparity ^23, 24^. For reference, at the viewing distance used (3.57 m), these are the correspondences between equivalent disparity and depth in units of physical distance (relative to the fixation point): -0.75 deg = -1.5 m, -0.25 deg = -0.69 m, -0.05 deg = -0.16 m, 0 deg = 0 cm, 0.05 deg = 0.18 m, 0.25 deg = 1.1 m, and 0.75 deg = 9.1 m. Because the object had a fixed physical boundary size, the greater depth range used in the monocular condition produced a greater range of image sizes. In the monocular condition, the retinal size of the object ranged from 0.23 deg, at the farthest depth, to 1.4 deg at the nearest depth. The five object velocities in world coordinates were 0, ±2.5, and ±5 cm/s, averaged over the duration of the Gaussian-velocity stimulus, and were the same in both binocular and monocular conditions. Self-motion velocities were also the same for binocular and monocular conditions, with average velocities (across trials) of ± 10cm/s (average velocity over the Gaussian-velocity stimulus profile). The average self-motion speed for each trial was chosen randomly from a Gaussian distribution centered around 10 cm/s with a standard deviation of 2.5 cm/s. We randomized both the direction and speed of self-motion to prevent subjects from associating a particular retinal image velocity with a particular relative depth.

### Control experiments: design and procedure

A total of six subjects were recruited for two control experiments. Subject 201 was one of the authors (GCD) and subject 225 also participated in the main experiment; the remaining 4 naïve subjects in the control experiments were different from the main experiment. Visual stimuli used in the control experiments were very similar to the main experiment, except as noted below.

#### Control Experiment 1: depth discrimination based on size cues only

The goal of this control experiment was to quantify how well subjects could judge object depth based only on size cues. The same visual scene was presented in this experiment; however, there was no simulated self-motion. The object remained stationary relative to the scene and was presented monocularly. Thus, the visual stimulus remained static over the course of 2 s, after which subjects made a choice about the object’s depth relative to the fixation point. Auditory feedback was given immediately after the choice was made. There were 90 trials in this control experiment (9 different depths x 10 repetitions). The only variable that changed from trial to trial was the object’s relative depth. The depth values used were 0, ±0.1, ±0.2, ±0.3, ±0.4 degrees.

#### Control Experiment 2: monocular detection of scene-relative object motion

The goal of this second control experiment was to quantify how well subjects could detect scene-relative object motion in the monocular viewing condition. The visual scene and self-motion profile were identical to the main experiment. After the stimulus disappeared, subjects had up to 1.5 s to indicate whether the object was moving or stationary relative to the scene by pressing one of two keys on a pre-labelled key pad (“not moving” vs “moving”). Auditory feedback was given immediately after the choice was made. The object’s depth was drawn randomly from a uniform distribution over the range from -0.45 to +0.45 degrees, and the object was presented monocularly. Object velocity in world coordinates took eleven different values: 0, ±1, ±2, ±3, ±4, and ±5 cm/s (averaged over the duration of the Gaussian-velocity stimulus). There were 20 repetitions for each of the non-zero object velocities and 100 repetitions for the zero object velocity. As the main experiment, there were two self-motion directions. Thus, there were a total of 600 trials in this control experiment. The resultant ratio of “not moving” to “moving” trials was 1:2 so that the subjects could not overly favor the “moving” choice.

### Behavioral data analysis

Subjects’ perceptual reports were converted to proportion of “far” choices for each unique combination of object speed in world coordinates and direction of object motion relative to self-motion, which we define as *signed object speed* (SOS). The magnitude of SOS is equal to the speed of object motion relative to the scene. The sign of SOS represents the direction of object motion relative to self-motion, where positive SOS means that object and observer move in opposite directions, and negative SOS means that object and observer move in the same direction. Because our predictions for depth biases (Figure 2) depend on the relative directions of object motion and self-motion, we sort the depth perception data by SOS. A psychometric function, *ψ*, based on a cumulative Gaussian function as the sigmoid, *S* (Equation 2), is fitted to data using psignifit 4.0 ^25^, where x is object depth (or object velocity for the second control experiment), *m* and *w* are the mean and standard deviation of the underlying Gaussian respectively, Φ is the standard cumulative Gaussian function, and *C* is a constant given by *C* = Φ^-1^(0.95) - Φ^-1^(0.05). The psychometric function also takes into account the guess rate, *γ*, and the lapse rate, *λ* (Equation 3).

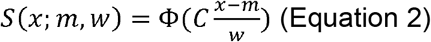

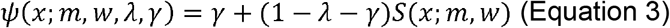

The point of subjective equality (PSE) is defined as the stimulus value at which the subject had an equal probability of choosing “near” vs “far” after accounting for the guess and lapse rates (such that the PSE does not necessarily occur at exactly 50% near/far choices).

### Eye movement analyses

During simulated lateral translation of the observer, the world-fixed fixation point translated horizontally on the display and subjects were asked to track the motion of the fixation point with their eyes. We first smoothed the fixation point and eye position data with a moving Gaussian window that served as a low-pass filter with cutoff frequencies of 10 and 15 Hz, respectively (function *filtfilt* in MATLAB). We used a lower cutoff frequency for the fixation point data because the position signal contained some high-frequency noise that was less prevalent in the eye signals. Although the movement of the fixation point was smooth, high-frequency noise was present in the signal about fixation point position because this signal was output over a digital-to-analogue converter and then sampled by the data acquisition system. Since self-motion speeds were randomized from trial to trial, we scaled the fixation point and eye position signals such that they would be expected to correspond to a fixed self-motion speed on each trial, and then we computed velocity profiles from the scaled position data. We removed saccades and blinks from the eye velocity data by finding local peaks (function *findpeaks* in MATLAB) in eye velocity that exceeded 10 deg/s, and replacing those portions of the velocity trace with the median eye velocity for that trial. Across subjects, there was an average delay of 80 ms between peak velocity of the fixation point and peak eye velocity. Thus, smooth pursuit gain was calculated by dividing the average eye velocity within a 200 ms window around peak eye velocity by the average angular velocity of the fixation point within a 200 ms window around its peak velocity. Calculation of pursuit gain depends only on the eye’s movement relative to the fixation point, and does not incorporate any information about the object’s velocity relative to the scene.

Pursuit gains were compared across conditions and subjects using one-way analysis of covariance (ANCOVA) models (function *aoctool* in MATLAB). To summarize pursuit gain at the group level, for each subject, we grouped every 3 repetitions of 70 conditions (7 different depths x 5 object velocities x 2 self-motion velocities) into a block of trials, such that there were 5 blocks of trials for each unique SOS. Therefore, the ANCOVA analysis was based on 150 data points (6 subjects x 5 blocks x 5 unique SOS values) when analyzing the influence of SOS on pursuit gain. When analyzing the influence of depth on pursuit gain, for each subject, we grouped every 3 repetitions of 50 conditions (5 different SOSs x 5 object velocities x 2 self-motion velocities) into a block of trials, such that there were 5 blocks for each unique depth. Thus, the ANCOVA analysis of the effect of depth on pursuit gain was based on 210 data points (6 subjects x 5 blocks x 7 unique depth values). We also performed linear regression on the data from each subject individually. For these individual analyses, the linear regression was based on 25 data points for SOS (5 blocks x 5 unique SOS values), or 35 data points for depth (5 blocks x 7 unique depth values).

### Model Fitting

To characterize the relationship between the magnitude of depth biases and SOS, the normalized point of subjective equality (PSE) was modelled as a polynomial function of *signed object speed* (SOS), as given by:

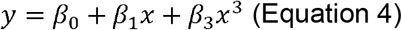

where *β*_0_, *β*_1_, *β*_2_ are the coefficients for each term, x is the signed object speed, and y is the normalized PSE. We obtained PSE values from the fitted psychometric functions for each subject, and normalized them within subjects by dividing by the maximum absolute value of PSE for each subject. For every subject, we grouped every 3 repetitions of 70 conditions (7 different depths x 5 object velocities x 2 self-motion velocities) into a block of trials, such that there were 5 blocks for each unique SOS. Therefore, for each subject, the fit of Eqn. 4 was based on 25 data points (5 blocks x 5 unique SOS values).

All linear regressions were performed using an iteratively reweighted least-squares algorithm to reduce the effect of outliers (Matlab function *fitlm* with *RobustOpts* turned on).

## Results

To test whether subjects can accurately judge depth from motion parallax in the presence of independent object motion, six naïve subjects performed a depth discrimination task in which they reported whether a target object appeared near or far relative to the fixation point during visually simulated self-translation (Figure 3). As the fixation point was rendered to be stationary in the virtual scene, subjects needed to generate smooth eye movements to maintain visual fixation during simulated self-motion (see Video 1 for example stimuli).

We first examine the depth biases associated with scene-relative object motion when subjects viewed the object monocularly. We next examine how the addition of binocular disparity cues influenced the biases induced by scene-relative object motion. We then investigate whether subjects’ pursuit eye movements may have played a role in their depth biases. Finally, we examine the shape of the dependence of depth bias on signed object speed (SOS).

### Scene-relative object motion strongly biases monocular depth judgements

Subjects’ reports were converted to proportion of “far” choices for each unique combination of object speed and direction of object motion relative to self-motion, which we define as *signed object speed* (SOS, see Methods). The sign of SOS is positive when object and observer move in opposite directions, and is negative when object and observer move in the same direction. We sort the depth psychometric data according to SOS because our predictions regarding depth biases depend on the relative directions of object motion and self-motion. A cumulative Gaussian function (Eqn. 3) was fitted to the proportion of “far” choices as a function of object depth. Point of subjective equality (PSE) is defined as the depth at which the subject had equal probability of reporting “near” vs “far”, after accounting for lapse rate and guess rate (see Methods). Horizontal error bars on psychometric curves indicate the 95% confidence interval of PSE.

Figure 4 shows data for an example subject, 233, tested in the monocular condition. In this condition, subjects can potentially judge the object’s depth using relative size and motion parallax cues. When the object moves in the same direction as self (negative SOS, Figure 4, purple lines), the subject shows a far bias that increases with object speed. This bias is consistent with our prediction if the subject bases their depth judgments on net retinal velocity and cannot isolate the component of object motion resulting from self-motion (Figure 2f). In contrast, when the object moves in the direction opposite to self-motion (positive SOS, Figure 4, blue lines), the subject shows a near bias, with greater bias for higher object speed. This is also consistent with our prediction (Figure 2f) that subjects report depth based on net retinal velocity.

**Figure 4:**
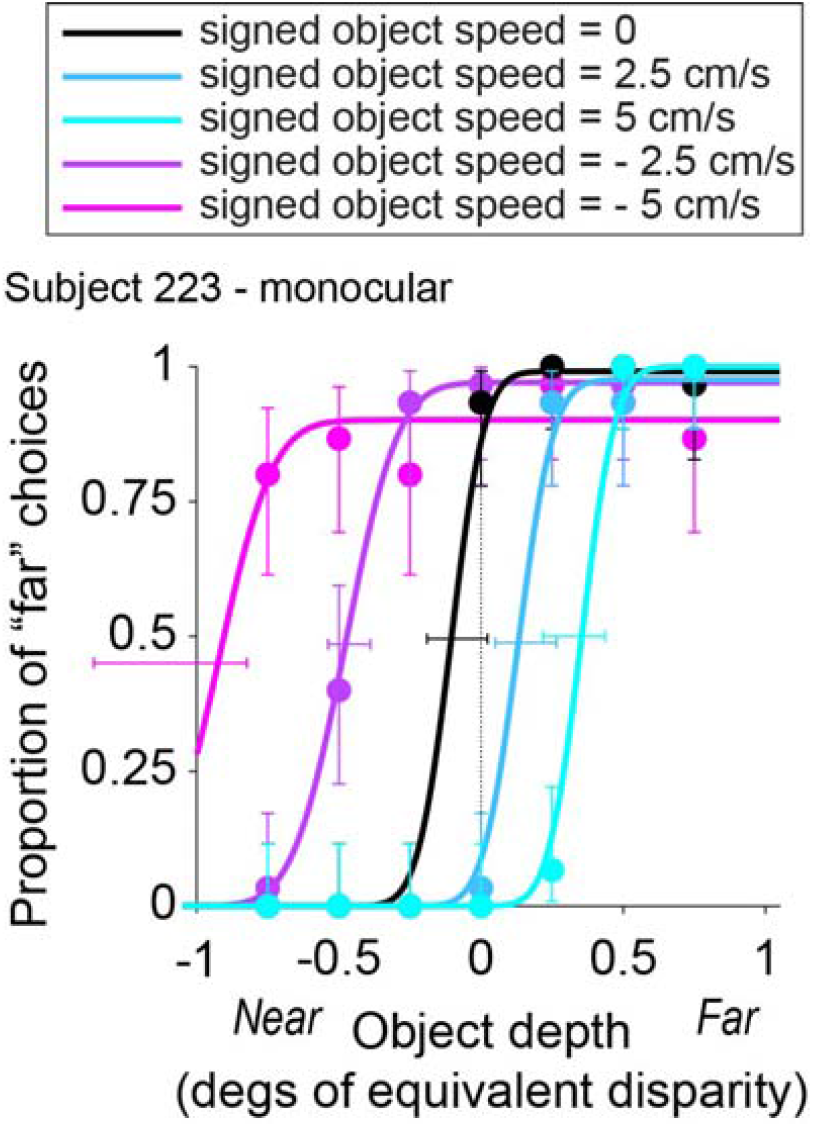
Data from an example subject in the monocular condition. Depth psychometric functions are shown for subject 223, color coded according to whether self-motion and object motion have the same direction (negative SOS, shades of purple), opposite directions (positive SOS, shades of blue), or there is no object motion (black). Different object speeds are indicated by color saturation. Vertical error bars indicate 95% confidence intervals for the proportion of “far” choices. Horizontal error bars indicate 95% confidence intervals for the PSE values obtained from the fits.

Indeed, all six subjects showed a similar pattern of results in the monocular condition (Figure 5a-f), with some variation in the magnitude of bias from subject to subject. To summarize and quantify these results for the monocular condition, we plotted PSE as a function of SOS for each subject (Figure 6a). For all subjects, PSE values increase monotonically with SOS. We performed a one-way ANCOVA with PSE as the dependent variable and with SOS and subject identity as independent variables. For the monocular condition, SOS (*F(1) = 273, p = 1*.*58×10*^-*34*^), subject identity (*F(5) = 10*.*2, p = 2*.*43×10*^*-8*^), and their interaction (*F(5) = 5*.*87, p = 5*.*94×10*^*-5*^) all show significant effects in predicting PSE values. When we perform linear regression for each subject separately (Bonferroni corrected significance level, *p = 0*.*00833*), we find that SOS has a significant effect on PSE for all six subjects: 222 *(t(23) = 8*.*97, p = 5*.*67×10*^*-9*^), 223 *(t(23) = 8*.*87, p = 7*.*00×10*^*-9*^), 224 *(t(23) = 9*.*54, p = 1*.*83×10*^*-9*^), 225 *(t(23) = 9*.*89, p = 9*.*44×10*^*-10*^), 226 *(t(23) = 18*.*6, p = 2*.*38×10*^*-15*^), and *229 (t(23) = 9*.*86, p = 9*.*96×10*^*-10*^).

**Figure 5:**
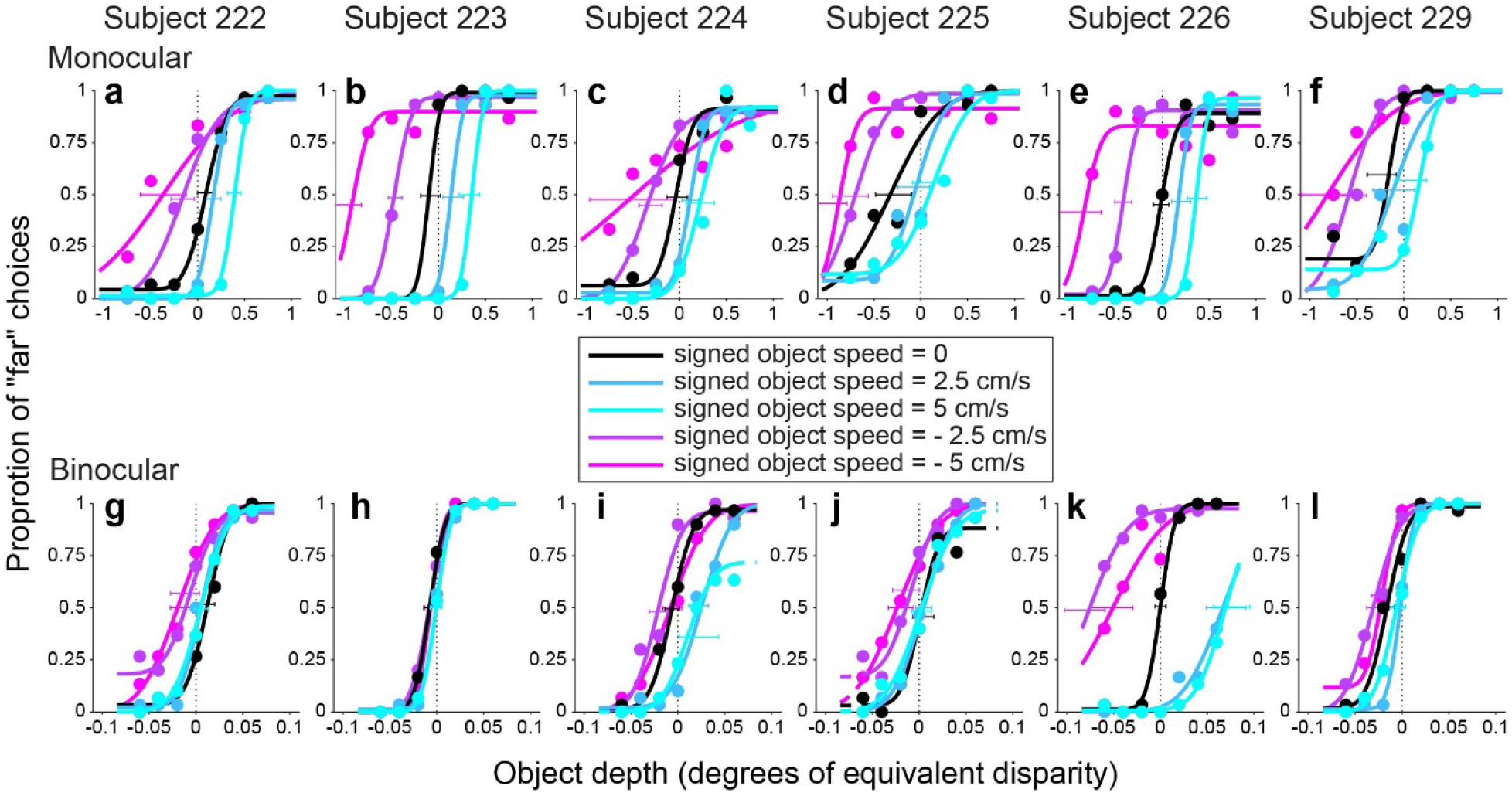
Behavioral results from all six subjects. a-f): Depth psychometric functions for the monocular condition from all six subjects. Note that the left end of the confidence intervals for SOS = -5 cm/s were cut off by the vertical axis in panels b, d, and e (for visual clarity). g-l): Depth psychometric functions in the binocular condition for the same subjects. Color scheme as in Figure 4.

**Figure 6:**
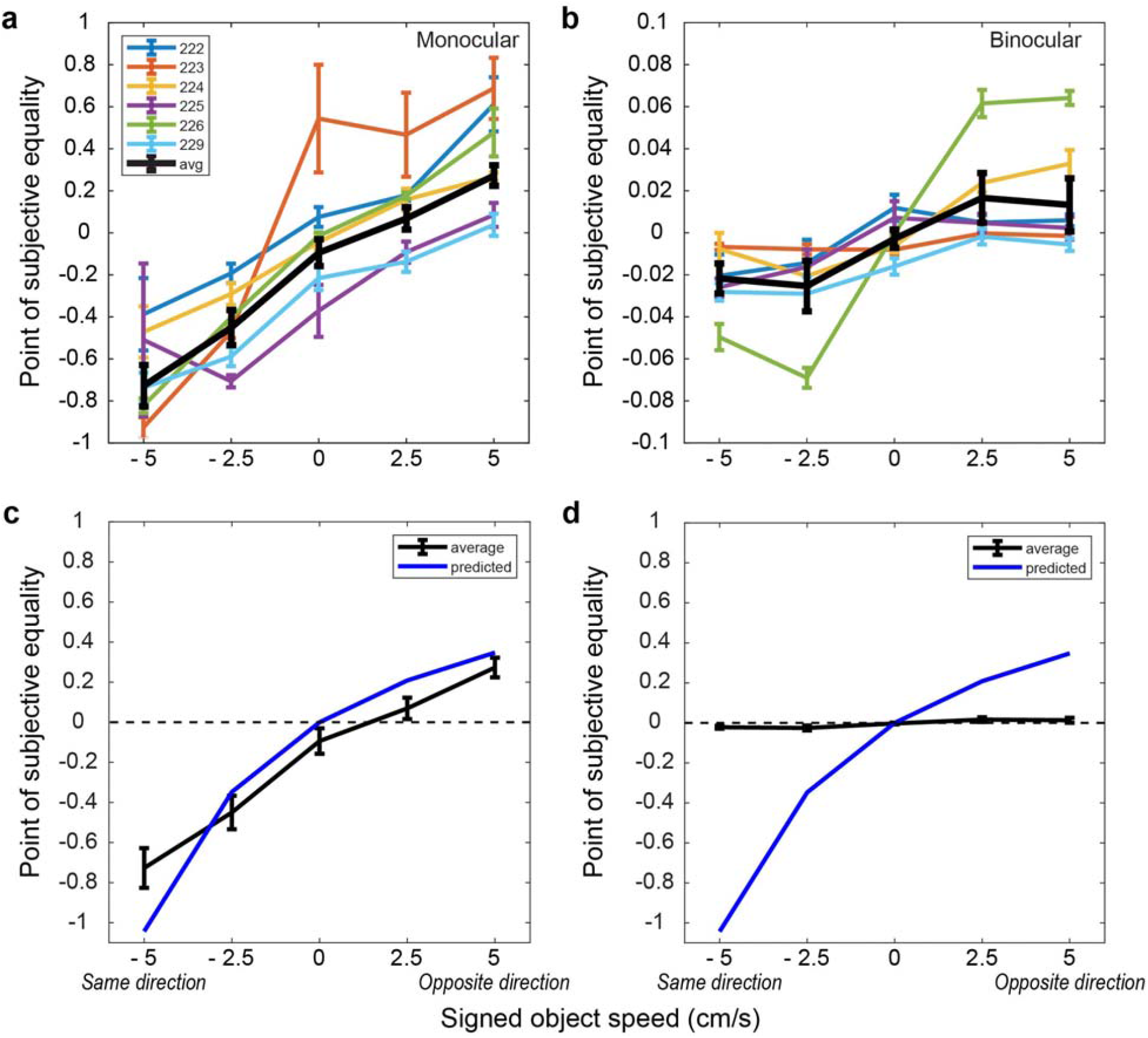
Summary of depth biases across subjects. a) PSE values for the monocular condition are plotted as a function of SOS for each individual subject (colored lines), as well as the group average (black). b) PSE values are shown for the binocular condition; format as in panel a. c) The group average data from the monocular condition (black) is compared with the predicted PSE values (blue) under the assumption that subjects estimate depth from the net retinal velocity of the object. d) Comparison between group average and predicted PSE values for the binocular condition; format as in panel c. Error bars for individual subjects represent SEM across blocks of trials; error bars for group averages represent SEM across subjects.

This significant positive effect in all subjects is consistent with our prediction that subjects do not compute depth from motion parallax by isolating the component of image motion related to self-motion, under the conditions of our experiments. To examine this further, we computed the predicted depth bias under the assumption that subjects compute depth based on net image motion. Figure 6c shows that the average biases of subjects in the monocular condition (black curve) closely follow this prediction (blue curve). The largest deviation from the prediction occurs when SOS = -5 cm/s, and this is likely caused by psychometric function fits being less well constrained for SOS = -5 cm/s due to the range of depth values tested (e.g., Figure 5b, d, e). Thus, in the monocular condition, our data show that subjects failed to parse out components of image motion caused by scene-relative object motion when judging depth, despite the presence of size cues.

### Reliability of depth cue modulates magnitude of depth bias

Perhaps the size cue was not sufficiently reliable to allow subjects to judge depth without confounding image motion resulting from self-motion and scene-relative object motion. If so, we might expect that perceptual biases induced by object motion would be eliminated, or at least greatly reduced, in the presence of binocular disparity cues. Thus, the same subjects were tested in a binocular condition in which disparity, motion parallax, and size could potentially all contribute to depth judgments. Note, however, that the size cue was weaker in the binocular condition since we used a much smaller depth range in this condition, owing to the greater sensitivity of binocular disparity cues.

As one might expect, depth judgements became much more accurate and precise in the binocular condition. Most subjects showed a similar pattern of biases—far bias when the object moves in the same direction as self and near bias when the object moves opposite to self—but the magnitude of biases (Figure 5g-l) was reduced by more than 10-fold compared with the monocular condition (Figure 5a-f). When we examine the relationship between PSE and SOS for the binocular condition (Figure 6b), we find the relationship to be less linear and more sigmoidal, as indicated by the group average (black curve). We further examine and quantify the shape of the relationship between PSE and SOS below.

We quantified the dependence of PSE on SOS and subject identity by performing a one way ANCOVA for the binocular condition. This reveals that SOS (*F(1) = 172, p = 5*.*65×10*^*-26*^), subject identity (*F(5) = 4*.*96, p = 3*.*29×10*^*-4*^), and their interaction (*F(5) = 28*.*9, p = 5*.*44×10*^*-20*^) all have significant contributions to predicting PSE values. When we perform linear regression for each subject separately, we find that SOS has a significant effect (*p < 0*.*00833*, Bonferroni corrected) on PSE in four subjects: 224 *(t(23) = 5*.*55, p = 1*.*20×10*^*-5*^), 225 *(t(23) = 3*.*55, p = 1. 69×10*^*-3*^), 226 *(t(23) = 9*.*65, p = 1*.*48×10*^*-9*^), and 229 *(t(23) = 4*.*20, p = 3*.*39×10*^*-4*^). For the remaining two subjects, there was no significant dependence of PSE on SOS after Bonferroni correction: 222 (*t(23) = 1*.*89, p = 0*.*0714*), 223 (*t(23) = 2*.*48, p = 0*.*0208*).

These results show that scene-relative object motion systematically biases depth judgements for most subjects, even in the presence of binocular disparity cues. However, it should be noted that these biases in the binocular condition are more than an order of magnitude smaller than both the biases observed in the monocular condition and the predicted biases based on computing depth from net retinal motion (Figure 6d).

### Smooth pursuit gain cannot explain depth biases caused by scene-relative object motion

Since the fixation point in our task is scene-fixed, subjects must smoothly rotate their eyes to maintain fixation during simulated self-motion. Stimuli are generated to simulated specific depths under the assumption that the eyes track the fixation point, and the analyses described so far assume that subjects accurately track the fixation point during simulated self-motion. Substantial systematic deviations in pursuit eye movements would alter the motion parallax presented and could potentially lead to altered depth perception of the stimulus. Could the depth biases that we have observed be explained by imperfect pursuit tracking? To assess this, we calculated pursuit gain (from eye and fixation point velocity traces) to quantify how well each subject tracked the fixation point (see Methods). The pursuit gain should be 1 when eye velocity was matched to velocity of the fixation point, less than 1 when subjects were under-pursuing, and greater than 1 when subject were over-pursuing. According to the motion-pursuit law (Equation 1), perceived depth should show a far (near) bias if subjects were under(over)-pursuing.

We extracted subjects’ eye velocity and computed pursuit gain for each SOS and depth condition for each subject. Figure 7a shows horizontal eye velocity traces for subject 223, tested in the monocular condition, for each SOS value (averaged across different depths). Since we did not enforce subjects to fixate before the stimulus started, the early deflections in the eye signal likely arise from the subject’s eyes settling on the fixation point at the beginning of the trial. For this subject, there are no significant differences in eye velocity between different SOS conditions (95% confidence intervals are overlapping), even though the subject shows large SOS-dependent depth biases (Figure 4).

**Figure 7:**
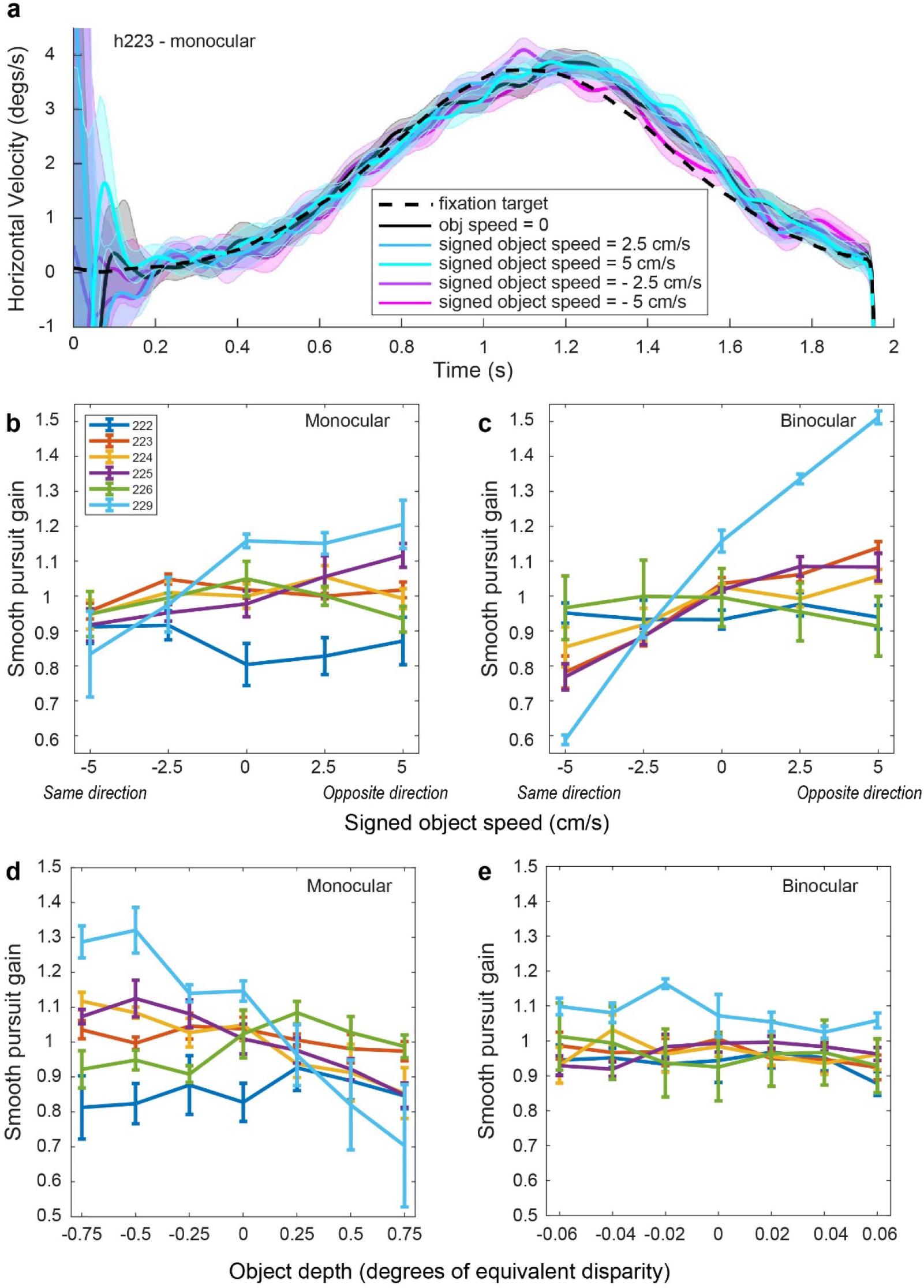
Example eye traces and pursuit gain analyses. a) Averaged eye velocity traces for different SOS conditions from subject 223 in the monocular condition. Shaded error bands represent 95% confidence intervals. b) Pursuit gain is plotted as a function of SOS for each subject (colored lines) in the monocular condition. Error bars represent SEM. c) Pursuit gain as a function of SOS for the binocular condition; format as in panel b. d) Pursuit gain plotted as a function of object depth for each subject in the monocular condition. Error bars represent SEM. e) Pursuit gain as a function of depth for the binocular condition; format as in panel d.

At the group level, pursuit gain does not show a consistent relationship with SOS in the monocular condition (Figure 7b). For subjects 225 and 229, the trend for pursuit gain to increase with SOS is consistent with the possibility that inaccurate pursuit might contribute to depth biases. However, for subjects 223, 224, and 226, pursuit gain essentially remains flat as a function of SOS, whereas pursuit gain tends to decrease with SOS for subject 222. We performed a one-way ANCOVA with pursuit gain as the dependent variable, and with SOS and subject identity as independent variables. For the monocular condition, pursuit gain depended significantly on SOS (*F(1) = 15*.*4, p = 1*.*38×10*^*-4*^), subject identity (*F(5) = 9*.*65, p = 6*.*38×10*^*-8*^), and their interaction (*F(5) = 7*.*22, p = 4*.*90×10*^*-6*^). At the individual subject level, SOS had a significant effect (*p < 0*.*00833*, Bonferroni corrected) on pursuit gain for only subjects 225 (*t(23) = 3*.*75, p = 1*.*03×10*^*-4*^) and *229 (t(23) = 3*.*33, p = 2*.*89×10*^*-3*^). For the binocular condition (Figure 7c), SOS (*F(1) = 154, p = 3*.*47×10*^*-24*^), subject identity (*F(5) =7*.*36, p = 3*.*83×10*^*-6*^), and their interaction (*F(5) = 36*.*9, p = 6*.*96×10*^*-24*^) again have significant effects on pursuit gain. Among individual subjects, SOS has a significant effect (*p < 0*.*00833*, Bonferroni corrected) on pursuit gain for subjects 223 (*t(23) = 9*.*34, p = 2*.*72×10*^*-9*^), 224 (*t(23) = 3*.*70, p = 1*.*19×10*^*-3*^), 225 *(t(23) = 8*.*04, p = 3*.*94×10*^*-8*^) and 229 (*t(23) = 24*.*4, p = 6*.*31×10*^*-18*^).

Although some subjects show a significant positive correlation between pursuit gain and SOS, it is unclear whether these effects are sufficiently large or consistent to account for the depth biases we have observed. To investigate this more directly, we performed a linear regression of the slope of the relationship between PSE shift and SOS against the slope of the relationship between pursuit gain and SOS, across subjects. We find no significant correlation between these slopes for the monocular (*R*^*2*^ *= 0*.*0228, F(4,6) = 0*.*0933, p = 0*.*775*) or binocular (*R*^*2*^ *= 0*.*229, F(4,6) = 1*.*19, p = 0*.*337)* conditions. Thus, we conclude that inaccurate pursuit cannot account for the depth biases induced by object motion. We also examined the Pearson’s correlation between SOS and each subject’s average vergence angle across depth values. We found no significant correlation in any of our six subjects *(p* > *0*.*0951)*, indicating that vergence angle did not systematically vary with scene-relative object velocity.

We also examined the relationships between pursuit gain and object depth for monocular and binocular conditions (Figure 7d and 7e). For the monocular condition, ANCOVA analysis revealed that depth (*F(1)= 43*.*7, p = 3*.*41×10*^*-10*^), subject identity (*F(5) = 10*.*0, p = 1*.*47×10*^*-8*^), and their interaction (*F(5) = 18*.*6, p = 4*.*03×10*^*-15*^) were significantly predictive of pursuit gain. When we performed linear regression for individual subjects, object depth had a significant effect (p < 0.00833, Bonferroni corrected) on pursuit gain for subjects 224 (*t(33) = -4*.*35, p = 1*.*24×10*^*-4*^), 225 (*t(33) = -5*.*33, p = 6*.*87×10*^*-6*^), and 229 (*t(33) = -5*.*69, p = 2*.*44×10*^*-6*^). For the binocular condition, only subject identity (*F(5) = 7*.*87, p = 8*.*73×10*^*-7*^) was significantly predictive of pursuit gain, and depth was not significantly correlated with pursuit gain for any of the subjects (*p > 0*.*0569*). We also examined the relationship between depth and vergence angle, and we found no significant correlation in any of our six subjects (Pearson’s correlation, *p > 0*.*429)*.

Together, these analyses indicate that neither pursuit gain nor vergence angle could account for our main findings regarding the biases in depth perception induced by scene-relative object motion.

### Quantitative examination of the shape between signed object speed and depth bias

The relationship between depth PSE and SOS was close to linear in the monocular condition (Figure 6a), whereas it showed a more sigmoidal dependence on SOS in the binocular condition (Figure 6b). To quantify this effect, we fitted linear regression models to normalized PSE values, for each subject individually, as well as the group average (Figure 8). Each linear model includes three terms: a constant, *SOS*, and *SOS*^3^ (see Methods). If PSE grows linearly with object speed, the *SOS*^3^ term would not be necessary to fit the data. However, if PSE saturates or declines with greater object speed, then *SOS*^3^ is more likely to be significant. For the monocular condition (Figure 8 a-f), none of the six subjects has a significant *SOS*^3^ term in the fit *(p > 0*.*418)*. For the binocular condition (Figure 8 h-m), subjects 224 (*t(22) = -2*.*21 p = 0*.*0378*), 226 (*t(22) = -8*.*34, p = 2*.*92×10*^*-8*^), and 229 (*t(22) = -2*.*16, p = 0*.*0418)* show a significant cubic term, *SOS*^3^ (Figure 8j, l, m).

**Figure 8:**
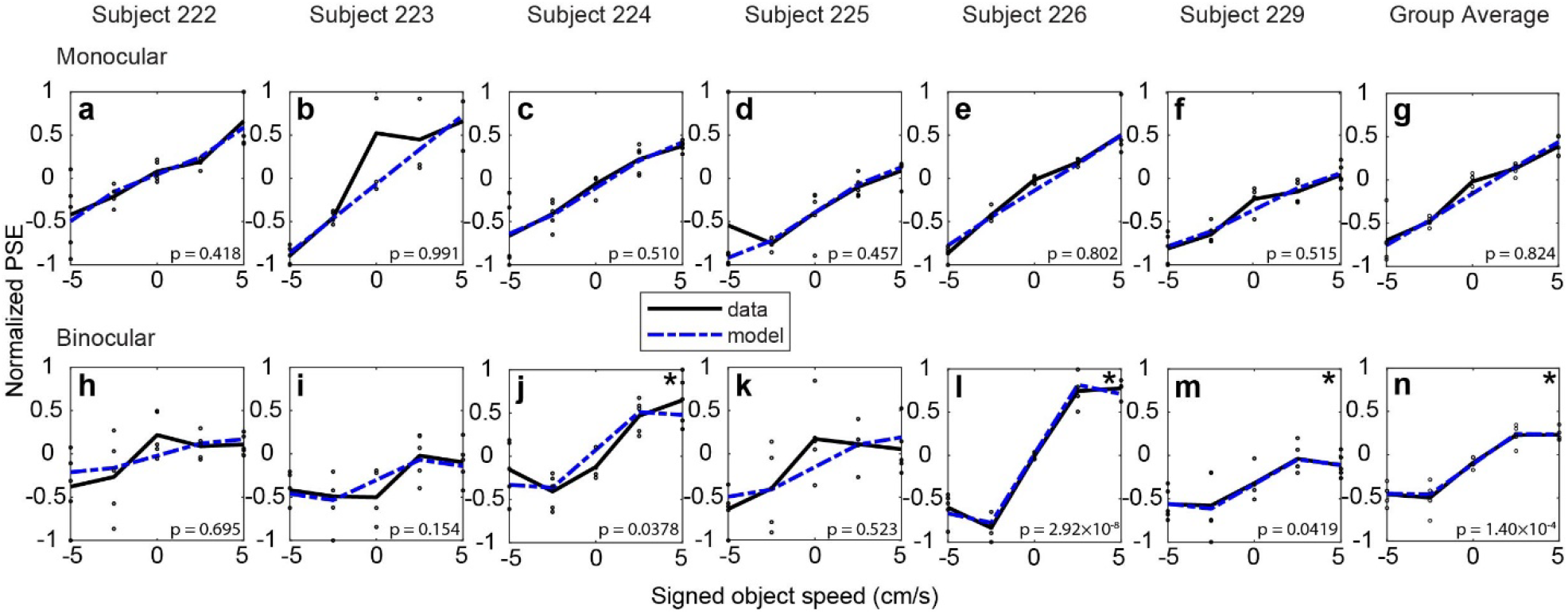
Quantifying the relationship between normalized depth PSEs and signed object speed. a-f): Normalized depth PSEs are plotted against SOS for each subject tested in the monocular condition. Each data point (black) represents the normalized PSE calculated by fitting a psychometric function to data from each block of trials (see Methods). Best-fit regression models are shown for each subject (dotted blue curves). g) Group average data and associated fit for the monocular condition; format as in panel a. h-m): Normalized PSE values are plotted against SOS for the binocular condition (black symbols), along with best fits of the regression model (dashed blue). n) Group average data and associated fit for the binocular condition. The p value shown in each panel indicates the significance of the *SOS*^*3*^ term in the model fit, which reflects the deviations from linearity. Significance of the *SOS*^*3*^ term is indicated by an * in the upper-right corner of some panels.

We also performed this analysis on the group average data for the monocular and binocular conditions (Figure 8g, n). At the group level, the cubic term *SOS*^3^ was significant for the binocular condition *(t(22) = -4*.*59, p = 1*.*40×10*^*-4*^) but not for the monocular condition *(t(22) = - 0*.*225, p = 0*.*824)*. This analysis demonstrates that the relationship between PSE and SOS is nonlinear for the binocular condition, and the potential implications of this finding are addressed in the Discussion.

### Size cues alone are sufficient to judge depth in the monocular condition

In the monocular condition, the depth biases induced by scene-relative object motion closely follow the prediction that is based on the assumption that subjects attribute all retinal image motion to self-motion (Figure 6c). This finding suggests that subjects placed very little weight on the size cues to depth in the monocular condition, and rather estimated depth from the net retinal image velocity while assuming that the object was stationary in the scene. One possible explanation for this finding is that the size cues in our stimuli were not sufficiently potent for subjects to estimate depth. To assess this possibility, in a separate control experiment, we asked a group of six subjects (including subject 225) to report the object’s depth using the same visual scene but without any scene-relative object motion or simulated self-motion. Thus, in this control experiment, size was the only depth cue available, since the object was presented monocularly.

Our results reveal that all subjects could indeed report depth rather well based on varying object size (Figure 9a-f). A logistic regression analysis reveals that all subjects showed a significant dependence of perceptual reports on depth specified by size cues (p<0.004 for each subject). Moreover, the average depth discrimination threshold (computed as the standard deviation of the cumulative Gaussian fit) in this size control experiment (mean ± SEM across subjects: 0.095 ± 0.0089) is somewhat lower than the average depth threshold of subjects (0.203 ± 0.0456) in the monocular condition of the main experiment, for the case where the object was stationary in the world (black curves in Fig. 5 a-f). While these mean values should not be compared quantitatively since mostly different subjects were used in the control experiment, this result strongly suggests that subjects *could* have used size cues to judge depth in the monocular condition of the main experiment. Yet, subjects apparently placed little weight on the size cues and mainly used motion parallax cues to judge depth in the monocular condition (see Discussion).

**Figure 9:**
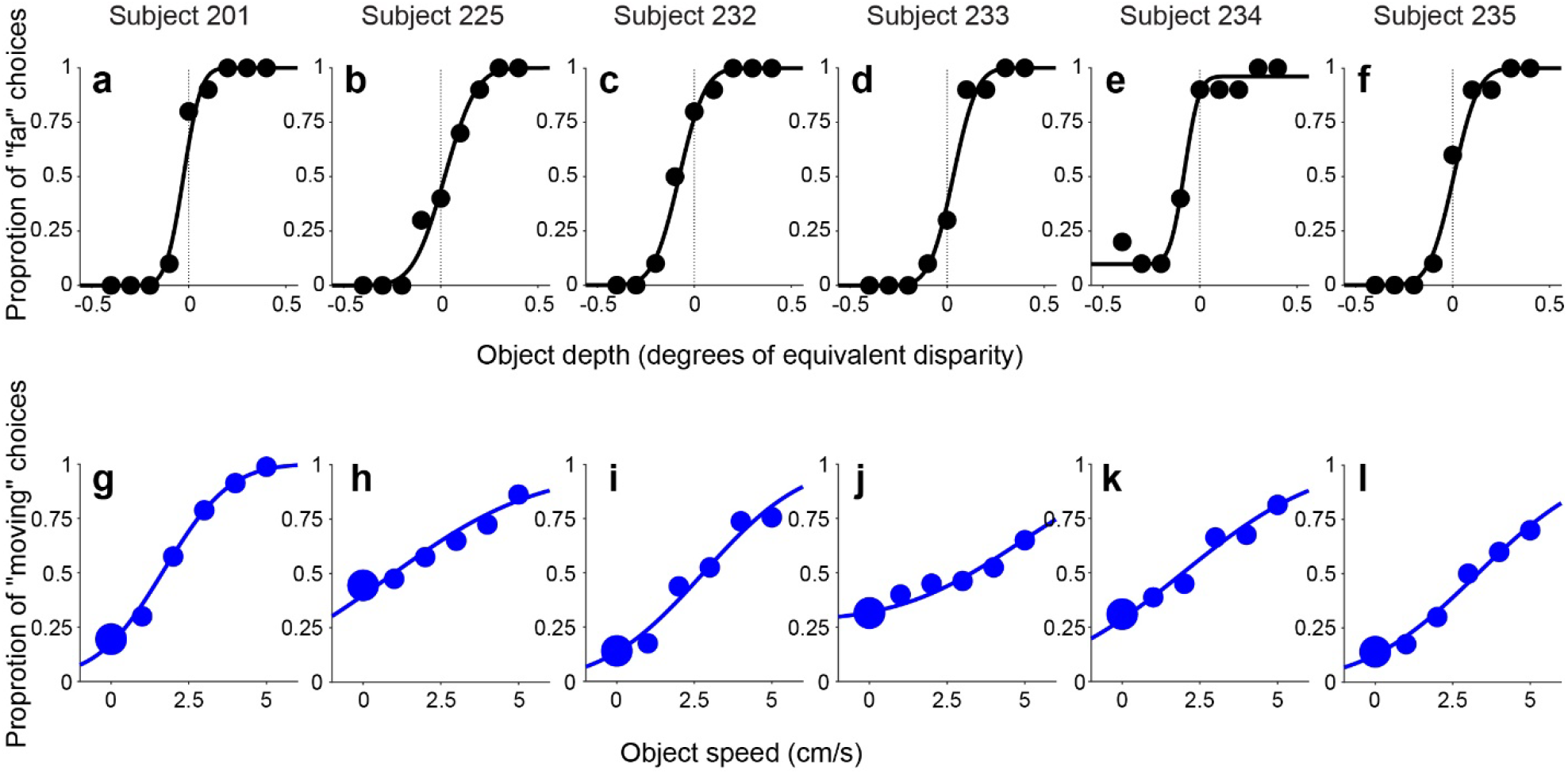
Subjects can judge depth and scene-relative motion of an object based on size cues. a-f) Depth psychometric functions for Control Experiment 1. Subjects can judge depth of a stationary object based only on size information when the object is presented monocularly. g-l) Psychometric functions for Control Experiment 2, in which subjects reported whether the object was moving or stationary relative to the scene. Proportion of “moving” choices is plotted as a function of the speed of the object’s motion relative to the scene (when speed = 0, the object is stationary in the scene). All subjects can detect object motion relative to the scene when the object is presented monocularly during simulated self-motion.

### Subjects are able to infer scene-relative object motion in the monocular condition

Another possible explanation for the result of Figure 6c is that subjects were not able to infer whether the object was moving or not relative to the scene in the monocular condition. If that were the case, then subjects would not be able to parse retinal motion into components related to object motion and self-motion, and it would make sense that their depth judgements were consistent with attributing all retinal image motion to self-motion (Fig. 6c). To address this possibility, in a second control experiment, we asked subjects to report whether the object moved relative to the scene, using the same visual display as the monocular condition of the main experiment. Comparable object velocities and depths were used in this control experiment (see Methods), and the object was again presented monocularly.

Our results demonstrate clearly that subjects are capable of inferring whether the object moved relative to the scene in our displays (Figure 9g-l) A logistic regression analysis demonstrated that all subjects showed a significant dependence of perceptual reports (“moving” vs. “not moving”) on object speed (p<0.00001 for each subject). Thus, subjects could indeed infer scene-relative object motion based on the size cues available in our monocular condition. Together, these control experiments demonstrate that depth biases seen in the monocular condition of the main experiment cannot be explained simply by subjects’ inability to use size cues to infer depth or object motion.

## Discussion

To accurately reconstruct a 3D scene, the visual system needs to compute the depth of the stationary components in the scene; in addition, the visual system needs to estimate depth for moving objects based on image motion that arises from either object motion or self-motion. We have shown that humans do not accurately judge the depth of moving objects during simulated self-translation, especially when static depth cues are less reliable as in the monocular condition.

The errors in depth judgements that we observe follow a systematic pattern: subjects show a far bias when the object and self are moving in the same direction, and a near bias when the object and self are moving in opposite directions. This pattern is expected if subjects interpret image motion as resulting from self-motion, and do not properly discount the component of image motion related to scene-relative object motion when computing depth. Perceptual biases were more than 10-fold greater when the object was presented monocularly than binocularly. However, depth biases were still significant in the binocular condition for most subjects. This suggests that, even in the presence of stereovision, subjects still place some weight on motion parallax cues to judge depth and they at least partially confound object motion in the world with motion parallax due to self-motion. Control experiments showed that size cues in the monocular condition are sufficient to report depth or judge object stationarity in the world, even though these size cues received little weight in the monocular condition of the main experiment, as discussed further below.

### Perceptual mechanisms underlying the motion-dependent depth bias

To understand the sources of this pattern of depth biases, we want to know the perceptual processes that might take place during such a task. We consider two ways that the brain might compute depth from motion parallax when objects move in the world. First, if the brain can successfully parse retinal image motion into components that are caused by object motion and self-motion ^26^, then depth could potentially be computed from the self-motion component using the motion-pursuit law ^12^. If the brain fails to attribute the correct amount of retinal motion to the object’s motion in the world, then the depth computation would be incorrect. Since previous literature has shown that flow parsing is often incomplete, having a gain less than unity ^27^, the component of image motion related to self-motion is likely to be misestimated, leading to erroneous depth estimates. However, if the flow-parsing gain is constant across stimulus conditions in our experiments, one might expect the PSE values in the binocular condition (Figure 6b) to have a linear relationship with SOS, but with a much flatter slope as compared with the monocular condition (Figure 6a). In contrast, we find a sigmoidal relationship between PSE and SOS for the binocular condition, and it is not clear how this might be explained solely by incomplete flow parsing.

Second, if a subject fails to recognize that an object is moving independently in the world, then the brain cannot correctly interpret the component of image motion caused by scene-relative object motion. Instead, the brain would have to explain away this extra component of image motion in terms of other variables, such as the object’s depth or the observer’s self-motion velocity. Consider a particular example (Figure 10): for an object that is nearer than the fixation point, leftward motion of the object in the world (Figure 10c) produces rightward image motion on the retina (Figure 10a, after accounting for image reversal by the lens). The same rightward image motion can be produced when the observer translates to the right and the object is stationary in the scene (Figure 10d), as well as by many other combinations of self-motion and object motion in the world (e.g., Figure 10e). If a subject is presented with the physical situation depicted in Figure 10e but they incorrectly infer that the object is stationary in the world, then they would have to explain away the component of image motion associated with the object’s motion. One possible perceptual reconciliation would be for the subject to believe that the object is nearer than it actual is, consistent with the physical situation depicted in Figure 10f. Thus, the subject’s (incorrect) belief that the object is stationary would lead to a near depth bias in this scenario. Hence, inference regarding the causes of image motion (causal inference) could also be a source of the systematic biases that we have observed.

**Figure 10:**
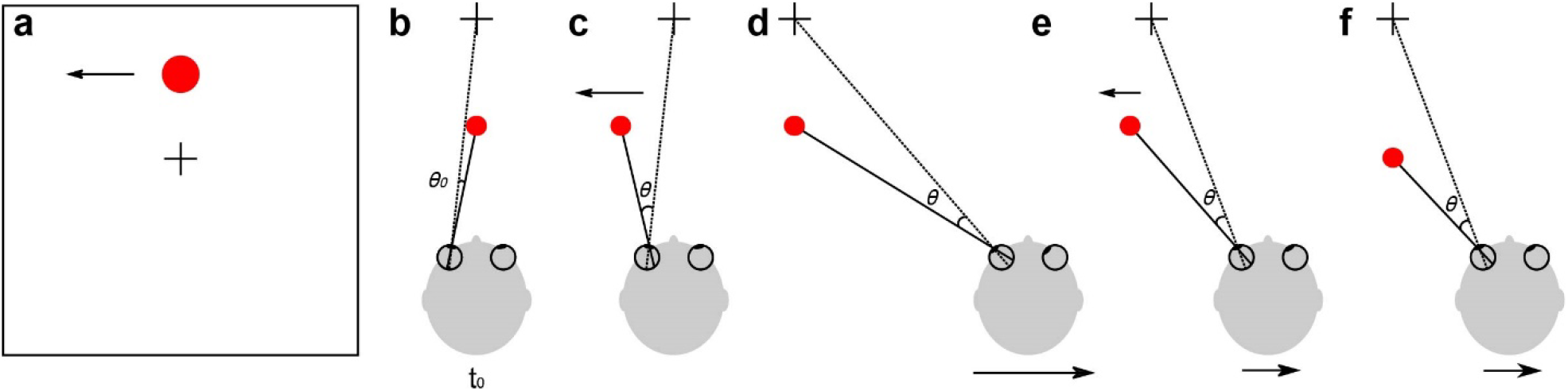
Multiple physical scenarios that could give rise to the same retinal image motion. a) Frontal view of the image motion (in screen coordinates) associated with an object (red circle). b-f) Schematic bird’s-eye view of a subject viewing the object in depth. b) The subject’s eyes are fixated on a world-fixed point (black cross). The starting positions of the fixation point and the object are at the midline (time t_0_). c-f): Physical situations that could all give rise to the same net image displacement of the object, ***θ***, at the end of the movement. c) The object moves leftward in the world while the observer is stationary. d) The observer moves rightward while the object is stationary in the world. e) Both object and observer move in the world. Many possible combinations of object and observer motion could produce the same image motion. f) The observer moves to the right more slowly than in d, but the stationary object is located nearer to the subject.

Although these two explanations (incomplete flow parsing and causal inference) are not mutually exclusive, the evidence from our binocular condition favors some contribution from causal inference. When subjects view the object binocularly, their perceptual biases do not increase linearly with greater object speed past 2.5 cm/s, leading to the sigmoidal shape of the group average data in Figure 6b. In contrast, biases continue to increase with object speed in the monocular condition, over the range tested (Figure 6a). We speculate that this difference is largely due to the greater reliability of depth cues in the binocular condition, such that subjects are less likely to believe that the object is stationary in the world when it is actually moving. Therefore, subjects have more information to correctly parse out the component of image motion related to self-motion and compute depth more accurately from that component. We observe a significant sigmoidal structure in the relationship between PSE and SOS for the binocular condition (Figure 8), which is consistent with predictions of a Bayesian causal inference framework and previous psychophysical observations that are taken as evidence for causal inference ^28-31^. Note that one might also expect a sigmoidal relationship for the monocular condition, in principle. However, because the monocular depth cues are less reliable, the range of object speeds that would be required to observe this effect is likely larger than the range sampled in our experiment. Additional experiments that involve judgments of both object motion and depth are ongoing to provide a more rigorous test of the causal inference explanation.

### Relative roles of size, motion parallax, and disparity cues

There were multiple depth cues available in our experiments. In the binocular condition, subjects could utilize binocular disparity and motion parallax cues, as well as relative size cues; in the monocular condition, there were relative size cues and motion parallax. However, the pattern of depth biases seen in the monocular condition is consistent with the prediction that subjects attributed most of the object’s retinal image motion to self-motion, and did not use size cues to discount object motion relative to the scene (Figure 6c). This was not because the size cues in the monocular condition were insufficient to judge depth, as we demonstrated in a control experiment (Figure 9a-f). Rather, in the monocular condition of the main experiment, subjects seemed to largely ignore the size cue and place much greater weight on motion parallax cues to depth. The reasons for this low weighting on size cues in the monocular condition are not clear. It is possible that subjects placed little weight on the size cues simply because they did not notice them and were not explicitly required to use them in the main experiment. Another possible, if not likely, explanation for the large depth biases in the monocular condition is that the direction and temporal profile of scene-relative object motion was compatible with self-motion. Under natural conditions, independently moving objects seldom move in a manner that can be easily attributed to self-motion. If an object moves vertically while the observer moves horizontally, for example, then scene-relative object motion is unlikely to be misinterpreted as being due to self-motion. Thus, our experimental design likely biased subjects toward interpreting image motion as resulting from self-motion in the monocular condition, despite the presence of robust size cues. Future studies should examine how adding additional cues to scene-relative object motion, such as directional or timing differences, alters the depth biases observed under monocular viewing conditions.

In the binocular condition, subjects were able to make use of the highly reliable binocular disparity cue to mostly discount the component of retinal image motion associated with object motion relative to the scene. This might suggest that the relative size cues in the monocular condition were not sufficiently potent to allow subjects to identify scene-relative object motion and discount it. Interestingly, however, when subjects were asked to report scene-relative object motion in a monocular control experiment, they were able to infer object motion from the combination of size and motion parallax cues (Figure 9g-l). It seems likely that the motion detection control experiment forced subjects to rely on size cues to parse image motion into components related to object motion and self-motion, as retinal velocity by itself offered no useful information about scene-relative object motion. These results suggest that task demands may substantially affect how subjects weight different cues to depth. In our experiments, subjects seemed to place little weight on size cues until they were required to do a (control) task for which utilizing size cues was essential. Thus, a useful topic for additional experiments would be to explore whether explicit training on size cues would alter depth biases seen in the monocular condition of the main experiment. It would also be useful to examine whether the modest biases observed in the binocular condition are eliminated under more natural conditions in which the timing and direction of scene-relative object motion is not fully consistent with self-motion.

### Pursuit signals in computing depth from motion parallax

In our experiments, self-motion was visually simulated and we did not incorporate physical translation that would generate vestibular signals. However, simulated self-motion combined with pursuit eye movements potentially provides sufficient information to compute depth from motion parallax. Previous studies have shown that it is the pursuit eye movement signal, not a vestibular signal that is used to compute depth from motion parallax ^24, 32^. The recorded eye data showed that subjects in our study were able to track the velocity of the fixation point throughout the trial with good accuracy (Figure 7a). Therefore, although there was no vestibular stimulation in the current experiment, subjects were able to judge depth sign from motion parallax.

### Future directions

The knowledge that the field has gained regarding computation of depth from motion parallax has been focused on stationary objects, including but not limited to how eye movements, head movements, or visual perspective cues affect computation of depth from motion ^16-18, 33, 34^, how motion parallax integrates with other depth cues to form depth percepts ^10, 20, 35^, and how other factors influence perceived depth from motion parallax ^15, 19, 21, 22^. Our work goes beyond the scope of current theory regarding computation of depth from motion parallax, namely the motion pursuit law ^12^ which applies directly only to stationary objects. Our work demonstrates that humans still put some weight on motion parallax in computing depth, even in the presence of more reliable disparity cues. Therefore, the biases that we observe in the binocular condition might affect vision in natural conditions, although this remains to be tested.

To tease apart the underlying perceptual processes for judging depth of moving objects, it would be beneficial to ask subjects to report whether they perceive the object to be moving in the world, in addition to reporting depth. We are currently conducting such a dual-report experiment, which may help to determine whether a causal inference process is involved in generating the observed biases. By conditioning on a subject’s report regarding scene-relative object motion, it should be possible to test whether perceived depth is influenced by causal inference.

Future neurophysiological studies based on this type of behavioral paradigm could also be valuable for understanding how the computation of depth for moving objects is carried out in the brain. Previous studies have shown that neurons in the middle temporal (MT) area can represent depth from binocular disparity ^36, 37^ and motion parallax ^23^ cues, at least for stationary objects. It is currently unknown how scene-relative object motion influences the neural representation of depth from disparity and motion parallax in MT, or whether the perceptual biases observed here might have a basis in MT population activity. Thus, exploring how the representations of depth and motion in area MT are modified by perception of scene-relative object motion may provide a good model system for studying key questions regarding the neural basis of causal inference.

## Acknowledgements

We thank Shelby Sabourin for help with behavioral data collection. This work was supported by an NEI F30 award to RLF (EY031183), and NIH grants to GCD (EY013644, U19NS118246), and by an NEI CORE grant (EY001319).

## Availability of Data and Materials

The datasets used and/or analysed during the current study are available from the corresponding author on reasonable request.

---

**Video 1: Example stimuli for the binocular condition**. Stimuli for two example trials are shown. In the first trial (same trial as illustrated in Figure 3) simulated self-motion is leftward, and the subject needs to make a smooth eye movement to the right to track the fixation point. The target object has a far depth, and a large leftward motion relative to the scene (5 cm/s). In the second trial, simulated self-motion is again leftward and the target object has a far depth, but there is no scene-relative object motion. In this second trial, motion of the target object relative to the fixation point is due to self-motion and the object’s depth. For correct viewing, the video should be viewed through red-green anaglyphic glasses, with the red filter placed over the left eye.

## Notes

### Competing Interest Statement

The authors have declared no competing interest.

